# Physical constraints on the positions and dimensions of the zebrafish swim bladder by surrounding bones

**DOI:** 10.1101/2024.07.04.602051

**Authors:** Koumi Satoh, Akiteru Maeno, Urara Adachi, Mizuki Ishizaka, Kazuya Yamada, Rina Koita, Hidemichi Nakazawa, Sae Oikawa, Renka Fujii, Hiroyuki Furudate, Akinori Kawamura

## Abstract

Precise regulation of organ size and position is crucial for optimal organ function. Since the swim bladder is primarily responsible for buoyancy in teleosts, early development and subsequent inflation of the swim bladder should be appropriately controlled with the body growth. However, the underlying mechanism remains unclear. In this study, we show that the size and position of the swim bladder are physically constrained by the surrounding bones in zebrafish. Non-invasive micro-CT scanning revealed that the anterior edge of the swim bladder is largely attached to the os suspensorium, which is an ossicle extending medioventrally from the 4th vertebra. Additionally, we observed that *hoxc6a* mutants, which lack the os suspensorium, exhibited an anterior projection of the swim bladder beyond the 4th vertebra. During the swim bladder development, we found that the counterclockwise rotation of the os suspensorium correlates with posterior regression of the swim bladder, suggesting that the os suspensorium pushes the swim bladder posteriorly into its proper position. Furthermore, our results revealed a close association between the posterior region of the swim bladder and the pleural ribs. In *hoxaa* cluster mutants with additional ribs, the swim bladder expanded posteriorly, accompanied by an enlarged body cavity. Taken together, our results demonstrate the importance of the surrounding bones in the robust regulation of swim bladder size and position in zebrafish.

## 1 Introduction

In animals, precise regulation of the size and position of internal organs is essential for their proper functioning (Rosello-Diez and Joyner, 2015; Texada et al., 2020). Disruptions in these regulatory mechanisms can result in genetic disorders and diseases. The early development of internal organs plays a crucial role in controlling their position and size. Once organs are formed, their dimensions are tightly controlled to synchronize with the growth of the body. Achieving allometric growth of organs involves regulatory mechanisms at the cellular level of individual cells composing the organs, which are further influenced by environmental factors. When it comes to regulation at the cellular level within organs, variables such as cell quantity, cell dimensions, and the quantities of extracellular components that form the organs assume pivotal roles in regulating organ size. In recent years, significant advancements have improved our understanding of the molecular mechanisms operating at the individual cellular level to control organ size (Dong et al., 2007; Gokhale and Shingleton, 2015; Penzo-Mendez and Stanger, 2015; Tumaneng et al., 2012). However, the role of external cues from the surrounding environments, such as the skeletal structures, in regulating organ size and position, especially concerning the internal organs of vertebrates, has been poorly understood.

To explore the regulation of the size and position of organs by the surrounding environments in vertebrates, the swim bladder, which some argue should rather be called the gas bladder in terms of function (Facey et al., 2022), may provide a good model. The swim bladder is a gas-filled sac located in the dorsal portion of the body that teleost fish uniquely acquired from the primitive lung (Facey et al., 2022; Funk et al., 2021; Funk et al., 2020). The primary role of the swim bladder in most teleost fishes is controlling buoyancy to maintain the vertical position in aquatic conditions, although it has specialized other roles in respiration, sound production, and sound reception in some species (Burton and Burton, 2018; Cook et al., 2024; Facey et al., 2022; Popper and Fay, 1993; Seymour et al., 2004). It is known that freshwater teleost fish tend to be larger than marine teleost fish in terms of the size of the swim bladder in their bodies because saltwater is denser and likely to produce more buoyancy (Facey et al., 2022). The swim bladder formation typically occurs at an early developmental stage in order to obtain buoyancy. In zebrafish living in freshwater, the swim bladder primordium initially emerges as a single lobe from the anterior portion of the primitive gut, followed by the subsequent formation of another anterior lobe at the anterior edge of the initial posterior lobe (Korzh et al., 2011; Parichy et al., 2009; Robertson et al., 2007; Wallace and Pack, 2003). After the formation of both the anterior and posterior lobes (SL 6.0-7.0 mm), the swim bladder undergoes inflation, which is closely correlated with the growing body, finally reaching up to 4-5 cm (Lindsey et al., 2010). During body growth, excessive inflation of the swim bladder leads to increased buoyancy, while poor inflation results in fish sinking due to decreased buoyancy. Therefore, maintaining the proper inflation of the swim bladder relative to body growth is crucial for fish survival. Failure in allometric growth in the swim bladder leads to energy loss in aqueous environments, which influences the effectiveness of essential teleost fish behaviors such as feeding, predator avoidance, and reproduction (Alexander, 1993). Thus, the precise regulation of swim bladder inflation with body mass throughout ontogeny remains elusive.

In this study using adult zebrafish, we developed a non-invasive X-ray micro-CT-based imaging method to visualize the swim bladder and the bones simultaneously. Our study reveals a close association between the swim bladder and the surrounding bones. Furthermore, our genetic study emphasizes the significance of neighboring bones in constraining both the size and position of the swim bladder. This leads us to propose that the scaling of the zebrafish swim bladder is physically regulated by neighboring bones during ontogeny.

## 2 Materials and methods

### Fish husbandry

Riken wild-type (RW) zebrafish were provided by the National BioResource Project Zebrafish in Japan and were maintained at 27°C with a 14 hour light/10 hour dark cycle. Embryos were obtained from natural spawning and the larvae were reared at 28.5°C. The stages of the larvae and juvenile fish were determined based on the standard length (SL), which corresponds to the distance from the snout to the caudal peduncle (Parichy et al., 2009). The allele of *hoxc6a* mutants used in this study is *sud122* (Maeno et al., 2024). The allele of *hoxaa* cluster-deleted mutant used in this study is *sud111* (Yamada et al., 2021). All the experiments using live zebrafish were approved by the Animal Care and Use Committee of Saitama University and conducted in accordance with the regulations.

### X-ray micro-computed tomography (CT) scanning analysis

X-ray micro-CT scanning was performed to simultaneously visualize the swim bladder and the surrounding bones of zebrafish. After the anesthesia with tricaine (MS222; Dojin), the adult zebrafish were sacrificed by immersion in ice water. The zebrafish were then subjected to micro-CT scanning without fixation. Since the zebrafish swim bladder is inflated with full air, the boundary between the lumen of the swim bladder and the air inside the swim bladder can be detected with the surrounding bones by using micro-CT scanning. We noticed that the fixation of adult zebrafish with 4 % paraformaldehyde (PFA) in PBS followed by the transfer to 70 % ethanol disrupted the morphology of the swim bladder. In addition, the boundary between the lumen of the swim bladder and the air within the swim bladder was barely detectable after such treatment. Using an X-ray micro-CT system (ScanXmate-CF110TSH320/460; Comscantecno) capable of imaging a wide field of view at high speed, each unfixed specimen was scanned at a tube voltage of 45 kV and a tube current of 70 μA and were rotated 360° in 0.18° steps to generate 2,000 projection images of 2,064 x 1,548 pixels. The CT scanning was performed in “continuous CT scanning mode” with a short scan time of approximately 3 minutes. The magnification during scanning of each specimen was determined for each sample according to the position and size of the swim bladder which can be confirmed by X-ray fluoroscopic images prior to scanning swim a result, each sample was scanned at a resolution of 6.5–8.0 µm/pixel. To reduce ta data volume, these micro-CT data were reconstructed at an isotropic resolution of 10.8–13.3 µm using the coneCTexpress software (Comscantecno) with a reduced resolution.

For scanning of the Weberian apparatus, adult zebrafish were analyzed by using an X-ray micro-CT (ScanXmate-E090S105; Comscantechno) as previously described (Akama et al., 2020; Yamada et al., 2021). 2D virtual sections and 3D images were constructed using OsiriX MD software (Pixmeo SARL) and Imaris v9.8 (Bitplane AG). Finally, movies were edited using Adobe Premiere Pro (Adobe).

### Genotyping of mutants

Genotyping of the larvae and mature fish was carried out by PCR. Genomic DNA was extracted from dissected tailfins by using the NaOH method (Meeker et al., 2007) and was used as a template for PCR. For the genotyping of *hoxc6a* frameshift-induced mutants, PCR was carried out as described (Maeno et al., 2024). For the genotyping of *hoxaa* cluster-deleted mutants, PCR was performed as described previously (Yamada et al., 2021). After the reactions, the PCR products were separated by the electrophoresis in 2 % agarose gel in 0.5 x TBE buffer or 15 % polyacrylamide gel in 0.5 x TBE buffer, and the genotype was determined.

### Visualization of skeletons in live zebrafish by calcein staining

To visualize the skeletons, calcein staining of zebrafish larvae was performed as previously described (Adachi et al., 2024; Akama et al., 2020). After several washings, the stained larvae were mounted in 2 % methylcellulose and photographed by a fluorescent stereomicroscope (Leica) or mounted in low-melting agarose gel and photographed by a confocal microscope (Olympus FV1000).

### Measurement of Feret diameter of the swim bladder and statistical analysis

Lateral images of the swim bladder with the adjacent bones obtained by micro-CT scan were compared. The size of the fish body was adjusted based on the length between the 1st and 9th rib-bearing vertebrae. The maximum Feret diameter of each anterior and posterior lobe was measured by ImageJ software. For statistical analysis, a student’s t-test was carried out.

## 3 Results

Including zebrafish, fishes of the series Otophysi (Cypriniformes, Characiformes, Siluriformes, and Gymnotiformes) of the superorder Ostariophysi comprise over 60% of freshwater fish species and exhibit the most acute auditory discrimination among living fishes with a wide range of auditory abilities (Facey et al., 2022; Nelson et al., 2016). This characteristic, which has evolved specifically in Otophysan fishes, is derived from a distinctive auditory system known as the Weberian apparatus (Bird and Hernandez, 2007; Diogo, 2009; Grande and Young, 2004; Maeno et al., 2024; Rosen and Greenwood, 1970). The Weberian apparatus consists of a chain of four small bones, namely, the scaphium, claustrum, intercalarium, and tripus, located in the anteriormost part of the vertebral column (Figure 2A-C). To enhance auditory sensitivity, these Weberian ossicles, connected via ligaments, move in response to the vibration of the swim bladder (similar but not homologous to three tiny bones of the mammalian inner ear) to transmit the vibrations of the swim bladder into fluid movements in the inner ears, providing the highest sensitivity and widest frequency range of hearing among the fishes (Facey et al., 2022). However, the precise nature of the interactions between the anterior portion of the swim bladder and the Weberian ossicles remains obscure. Therefore, we developed high-resolution imaging using micro-CT scanning, without any chemical fixation and invasion, to precisely compare the relationship between the swim bladder and the surrounding bones in zebrafish. In fixed specimens, it has been possible to visualize the Weberian apparatus and the swim bladder independently (Yamada et al., 2021), but simultaneous visualization has proved challenging. We hypothesized that if a gas-filled bladder were replaced with a liquid such as a fixative solution, the X-ray transmission would not be different, thus obscuring the boundary between the swim bladder and the surrounding cells. Therefore, we used unfixed specimens and found that the luminal boundaries of the swim bladder could be concurrently detected with the surrounding bones because the X-ray transmission rates are sufficiently different between the swim within the bladder and the surrounding cells. Using this method in wild-type adult zebrafish, we found that the anterior portion of the anterior lobe is attached to two different types of bones: the posterior edge of the tripus (fan-shaped Weberian ossicle on the 3rd vertebral body, centrum) and the os suspensorium (bone extending mid-ventrally from the 4th centrum) (Figure 1A). The tripus is an ossicle in the Weberian apparatus whose posterior end contacts the swim bladder and is thought to be the first bone to transmit swim bladder vibrations. On the other hand, os suspensorium exhibits morphological diversity even among otophysan fishes, and some extinct primitive otophysan fishes did not possess os suspensorium (Bird and Hernandez, 2007; Britz et al., 2021; Diogo, 2009), therefore the os suspensorium is generally not included in the Weberian apparatus and its role remains unknown. While the connections between these bones and the swim bladder have been previously described in zebrafish (Bird and Mabee, 2003; Bird et al., 2020; Grande and Young, 2004), our 2D micro-CT sections clearly showed that a substantial portion of the os suspensorium physically contacted with the anterior lobe because it appears no gap between the surface of the swim bladder and the os suspensorium (Figure 1B, C). In addition, the surface of the os suspensorium facing the swim bladder appears curved to follow the surface of the swim bladder smoothly, whereas the opposite side is apparently not (Figure 1F, G). Furthermore, the contact point between the os suspensorium and the swim bladder is observed at the front edge of the swim bladder, whereas the point of contact between the tripus and the swim bladder is located more posteriorly (Figure 1B-E). Based on these observations, our results suggest that the zebrafish os suspensorium interacts more physically with the anterior end of the swim bladder compared to tripus.

**FIGURE 1.**
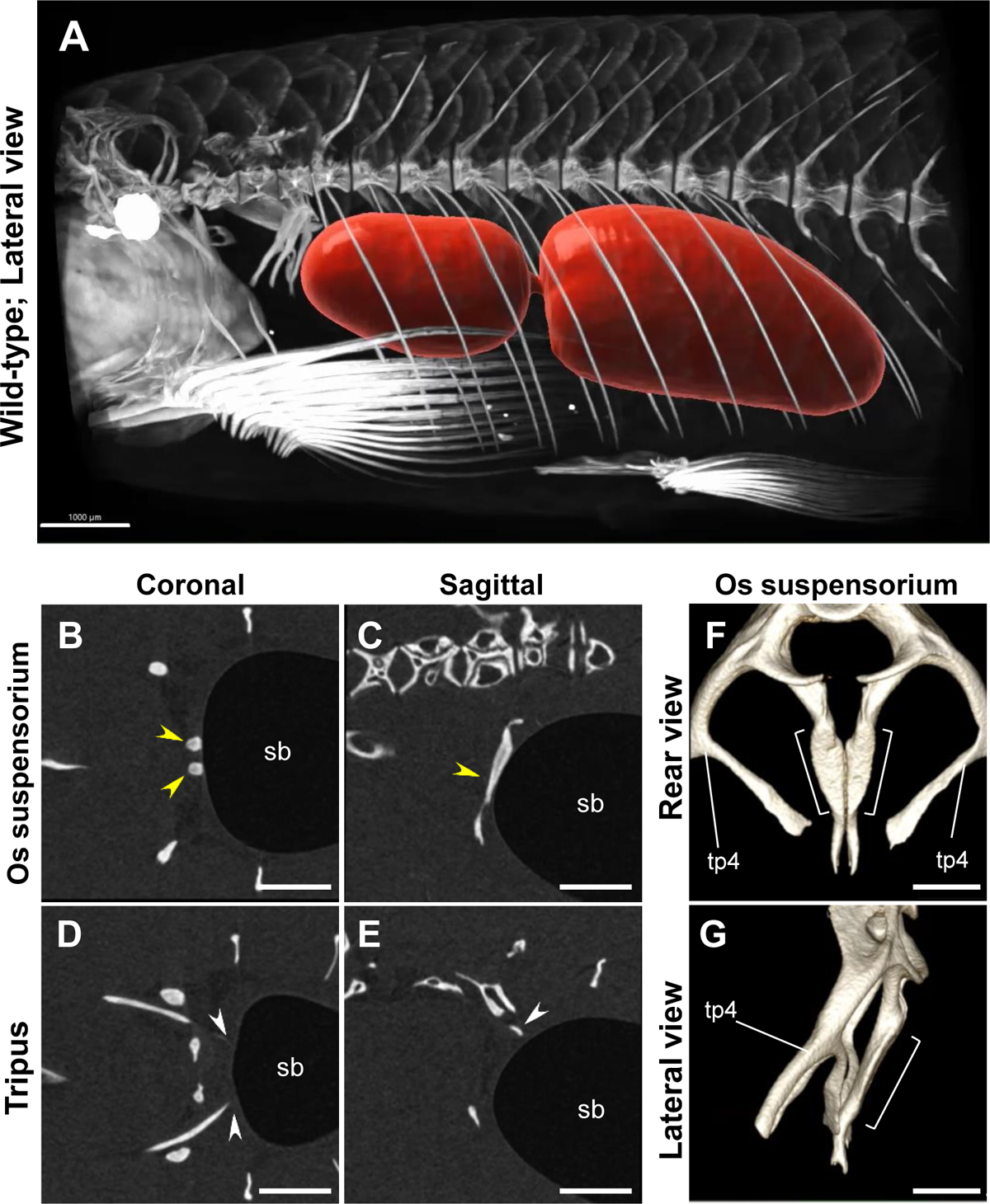
The os suspensorium is more in contact with the anterior end of the swim bladder than the tripus in adult zebrafish. (A) A micro-CT scan image demonstrates the relationship between the swim bladder (red) and the surrounding skeletal structures (white) in wild-type adult zebrafish (*n*=4). (B-E) Plane sections obtained through micro-CT scanning. The yellow arrowhead indicates the os suspensorium, which directly attaches to the anterior margin of the swim bladder (sb). The white arrowhead represents the posterior edge of the tripus, which is closely associated with the swim bladder. The anterior orientation is to the left. (F, G) The morphology of the os suspensorium is shown. The os suspensorium is an extending bone located medially from the 4th centrum. The portion of the os suspensorium in contact with the swim bladder is demarcated by a bracket. The transverse process of vertebra 4 (tp4), which extends laterally and ventrally from the 4th centrum, is also indicated. Scale bars: 1000 μm (A), 500 μm (B-E), 300 μm (F, G).

**FIGURE 2.**
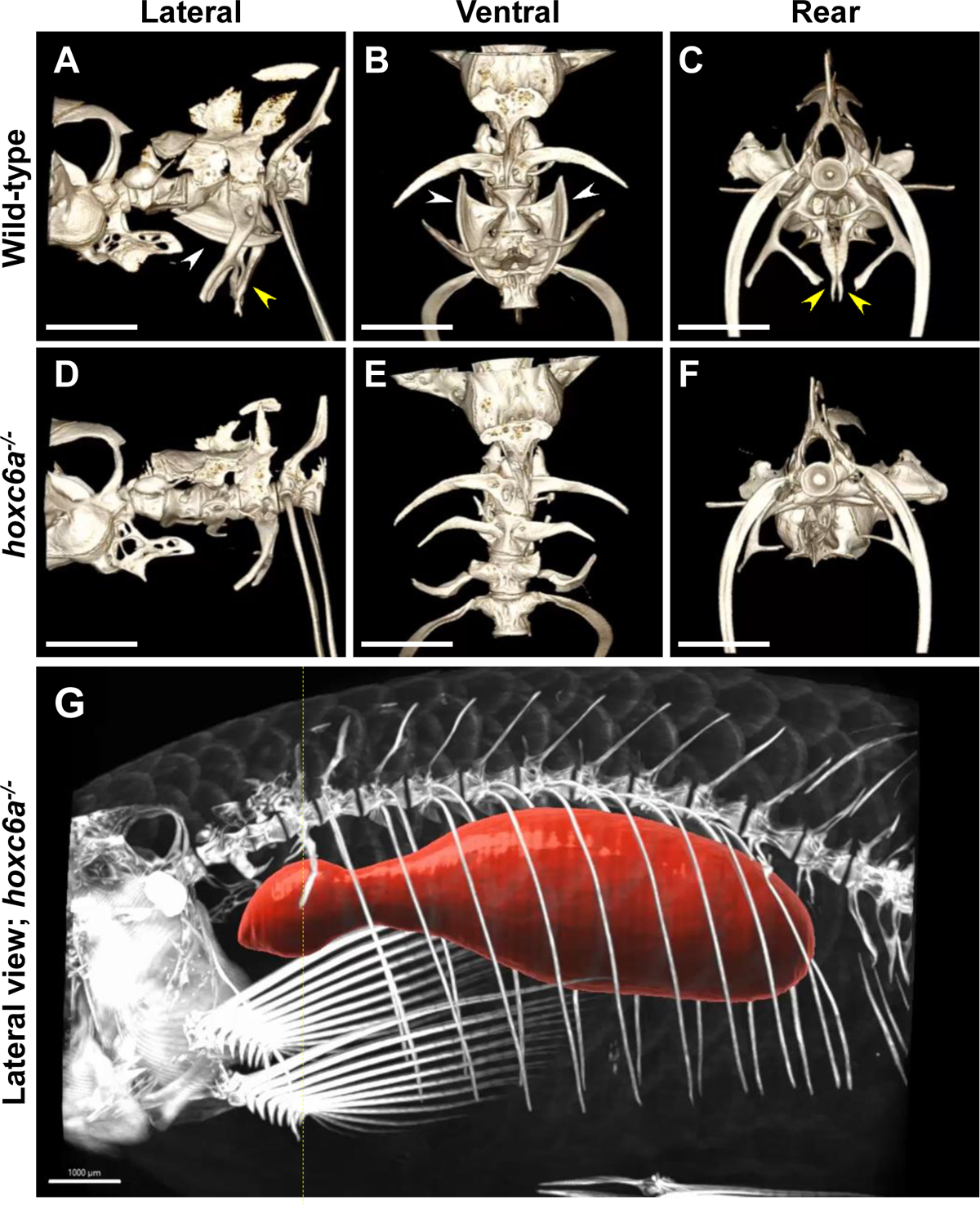
Anterior protrusion of the swim bladder in *hoxc6a* mutant fish showing the absence of the os suspensorium and the malformed tripus. (A-F) Representative micro-CT scan images of the anterior vertebrae in wild-type (*n*=4) and *hoxc6a* homozygous adult fish (*n*=4). In the images, the os suspensorium is indicated by the yellow arrowhead, while the tripus is indicated by the white arrowhead in wild-type fish. (G) Visualization of the swim bladder (depicted in red) and the surrounding bones in *hoxc6a* mutant fish (*n*=4). The presumptive anterior edge of the swim bladder in normal adult fish is indicated by the yellow dashed line. Scale bars: 1000 μm.

To examine the function of the os suspensorium in the swim bladder formation, we focused on *hoxc6a* mutants. In our previous study, zebrafish *hoxc6a* mutants exhibited the absence of os suspensorium on the 4 centrum, and the tripus on the 3rd centrum was transformed into the lateral process-like bones on the 2nd centrum (Maeno et al., 2024). We also found defects in the formation of the swim bladder in *hoxc6a* mutants (Supplementary Figure S1). In *hoxc6a* larvae, the swim bladder phenotype could be classified into two cases: one with a single, enlarged swim bladder as was observed in *hoxca* cluster mutants (Yamada et al., 2021), probably due to the improper formation of the anterior lobe. The other is where the anterior lobe emerges after the posterior lobe forms but with relatively inadequate inflation. One possible explanation for this phenotypic variation is that other *hox* genes in *hoxca* cluster, which are co-expressed in the foregut where the swim bladder originates (Zheng et al., 2011), compensate to some extent for the *hoxc6a* function. To investigate the role of the os suspensorium, we visualize the swim bladder and the surrounding bones in *hoxc6a* mutant fish by micro-CT scanning. In wild-type fish, the anterior end of the swim bladder is positioned below the 4th centrum in contact with the os suspensorium (Figure 1A). However, in *hoxc6a* adult mutants, possessing both the anterior and posterior lobes of the swim bladder, we observed that the anterior edge of the anterior lobe extended more anteriorly beyond the 4th centrum while showing abnormal morphology of the swim bladder (Figure 2G). These results suggest an interesting possibility that the os suspensorium serves as a physical barrier, determining the correct positioning of the anterior edge of the swim bladder in zebrafish.

To gain further insights into how the os suspensorium terminates the anterior edge of the swim bladder, we analyzed the development of the swim bladder in more detail. In zebrafish around the SL 6.0 mm, the anterior lobe is known to emerge as a small bulge from the posterior lobe, followed by rapid inflation of the anterior lobe (Parichy et al., 2009). Intriguingly, we found that the anterior end of the swim bladder expanded to reach near the 2nd centrum in wild-type juvenile zebrafish (Figure 3A*-*A*’’*). This implies that the anterior end of the swim bladder advanced beyond the 4th centrum, the typical position of the anterior end of the swim bladder in normal adult zebrafish (Figure 1A). Subsequently, we observed that the anterior end of the swim bladder gradually retracted to a location near the 4th centrum (Figure 3B*-*B*’’*). In contrast, in *hoxc6a* mutants with inflated anterior lobe, the swim bladder extended to around the 2nd centrum, similar to the wild type (Figure 3C*-*C*’’*), but the anterior lobe failed to recede as was observed in wild-type and remained anteriorly (Figure 3D*-*D*’’*).

**FIGURE 3.**
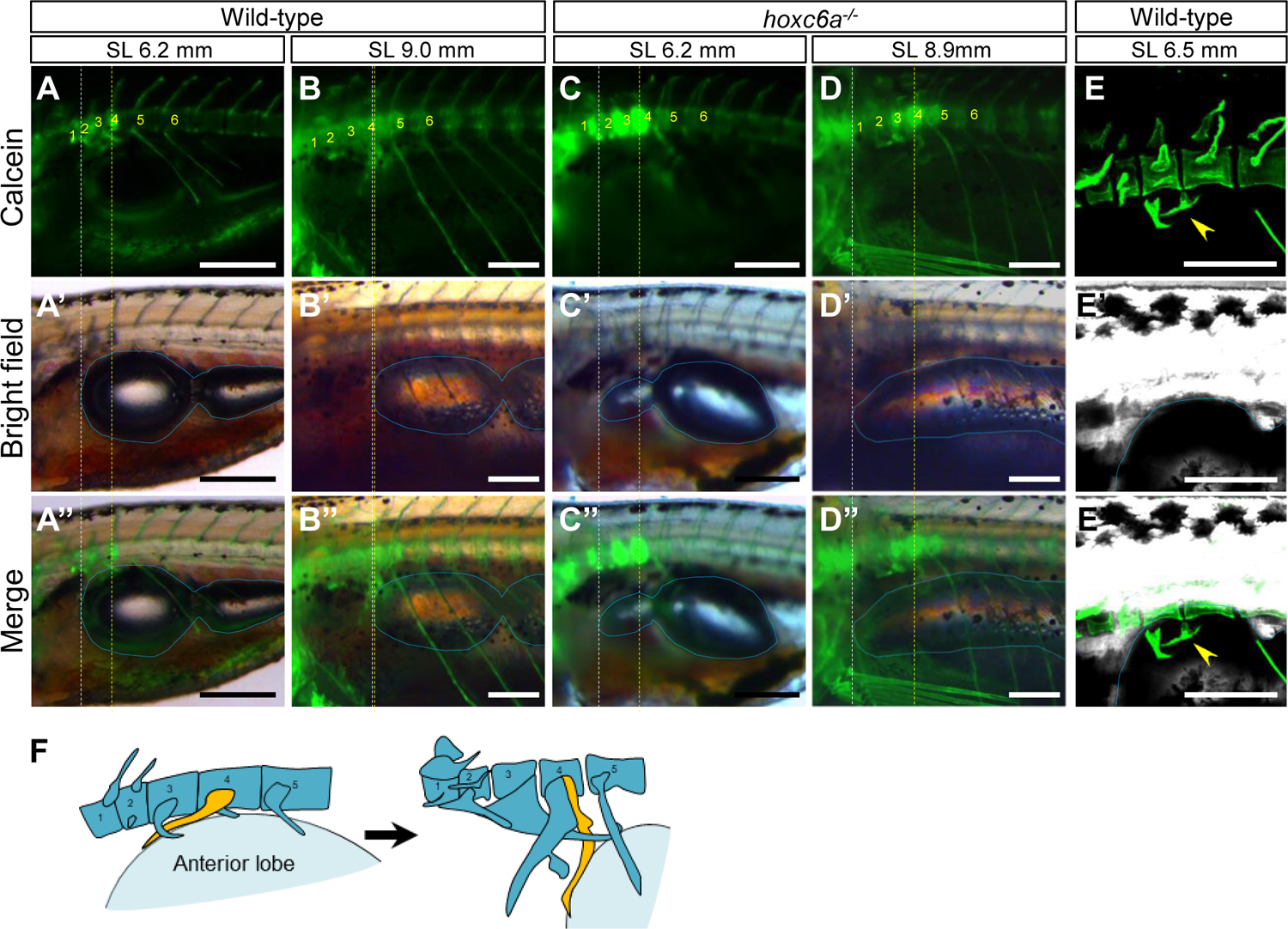
Pushback of the anterior lobe of the swim bladder by the developing os suspensorium. (A-D) The anterior edge of the inflating swim bladder relative to the vertebrae was compared between wild-type (*n*=6) and *hoxc6a* mutants (*n*=5). Calcein staining was used to visualize the bones. The anterior edge of the anterior lobe in the swim bladder is highlighted with a white dashed line. The presumptive position (4th centrum) corresponding to the anterior edge of the swim bladder in typical normal adult zebrafish is indicated by a yellow dashed line. The swim bladder is outlined by a blue dashed line. The number of vertebrae starts from the most anterior vertebra. Images were taken using a fluorescent stereomicroscope from the lateral side. (E) After the calcein staining, the anterior vertebrae and swim bladder of wild-type larvae were visualized using a confocal microscope (*n*=3). An oblique dorsoventral view is presented, and the arrowhead indicates the developing os suspensorium attaching to the swim bladder. (F) Schematics illustrate the model showing the counterclockwise rotation of the os suspensorium (depicted in yellow) pushing back the anterior swim bladder (light blue) during zebrafish development. Scale bars: 300 μm.

Concurrently, the development of the os suspensorium has been described to be characteristic in zebrafish (Bird and Mabee, 2003; Grande and Young, 2004). The os suspensorium initially emerged as an anteriorly elongating bone from the basiventral region of the 4th centrum and gradually rotated towards the ventral position, where the os suspensorium is situated in adult fish. Significantly, the period during which a counterclockwise rotation of the os suspensorium occurred coincided with the developmental stage of the anterior lobe inflation. Therefore, we compared the development of the os suspensorium with that of the anterior lobe of the swim bladder. In wild-type larvae at approximately SL 6.5 mm, we observed that the os suspensorium seemed to attach to the inflating anterior lobe (Figure 3E*-*E*’’*). Consistent with this model, the retraction of the inflated anterior lobe was not markedly evident in *hoxc6a* mutants lacking the os suspensorium. These findings suggest that the anterior lobe of the wild-type swim bladder becomes excessively inflated and is subsequently pushed posteriorly due to the mechanical force driven by rotation of the os suspensorium, eventually ensuring its proper ventral positioning ventral to the 4th centrum (Figure 3F). Furthermore, our results suggest that the anterior protrusion of the anterior lobe in *hoxc6a* mutants is due to the lack of pushback by the os suspensorium, thereby maintaining the anteriorly protruded position that initially occurs during development.

Our results revealed the precise termination of the anterior lobe by the os suspensorium. These findings give rise to a further question of how the posterior region of the swim bladder is regulated in zebrafish. After the anterior front of the anterior lobe is fixed below the 4th centrum, we observed that the posterior chamber in zebrafish undergoes continued expansion in the posterior direction, consistent with the previous observation (Parichy et al., 2009). A micro-CT scan analysis of wild-type adult fish has unveiled a close association between the posterior lobe of the swim bladder and the pleural ribs, particularly the most posterior two pairs of the ribs (Figure 1A), whereas the anterior lobe does not exhibit such an association with neighboring pleural ribs.

These observations prompted us to investigate whether the position and dimensions of the swim bladder are influenced in zebrafish possessing additional pairs of pleural ribs. In our previous study, *hoxaa* cluster-deficient fish were found to possess 10 pairs of ribs, in contrast to the typical 9 pairs in wild-type zebrafish (Yamada et al., 2021). Using micro-CT scanning on *hoxaa* cluster mutants, we observed that the posterior lobe of the swim bladder extended further posteriorly in conjunction with the additional pairs of ribs (Figure 4A*-*F). The expansion occurs in the posterior lobe, while the anterior lobe of the swim bladder does not show a noticeable change in size. Consistent with these observations, we found that the maximum Feret diameter of the posterior lobe significantly increased in *hoxaa* cluster mutants compared with that in wild-type fish, while the maximum Feret diameter of the anterior lobe was not significantly changed (Figure 4G). These results suggest that the inflation of the posterior lobe is constrained by the cavity enclosed by the pleural ribs.

**FIGURE 4.**
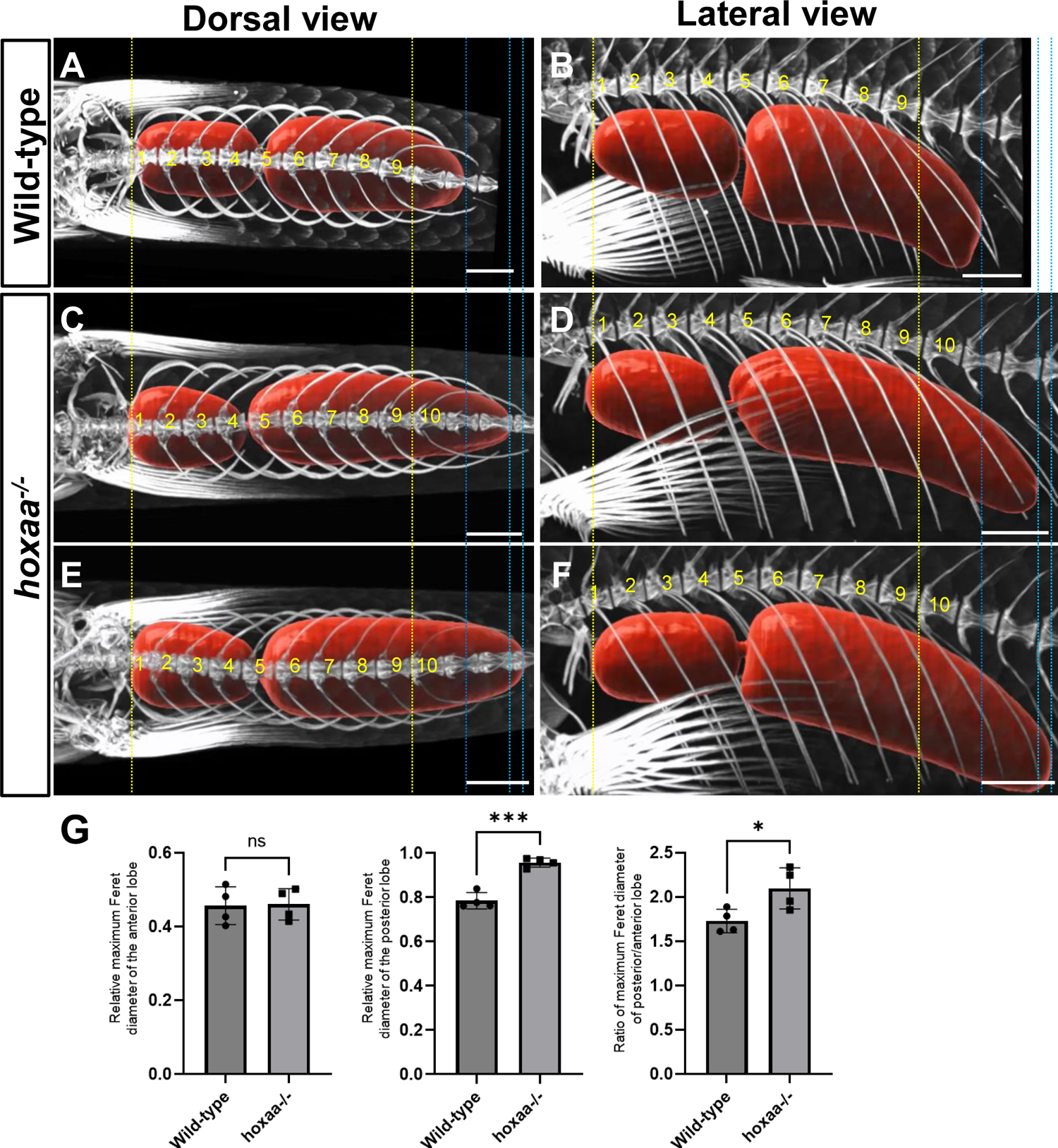
Posterior enlargement of the swim bladder in *hoxaa* cluster-deleted mutants with an additional pair of pleural ribs. (A-F) Micro-CT scan images showing the swim bladder (red) and the surrounding skeletal structures in wild-type (*n*=4) and *hoxaa* cluster-deleted adult fish (*n*=4). To ensure reproducibility, two representative images are shown for *hoxaa* cluster-deleted adult fish. The vertebrae are numbered in order starting from the first rib-bearing vertebra. To facilitate comparisons, the dimensions of the fish body were standardized based on the distance between the first and ninth rib-bearing vertebra, shown by the yellow dashed line. The blue dashed line indicates the posterior extent of the swim bladder in wild-type fish, while the light blue dashed lines represent the posterior extent of the swim bladder in *hoxaa* cluster-deficient mutants. (G) A comparison of the maximum Feret diameter of the anterior and posterior lobes between wild-type and *hoxaa* cluster-deficient mutants. A Student’s t-test was performed, with significance indicated as *PL<L0.05 and ***PL<L0.001. Scale bars: 1000 μm.

## 4 Discussion

In this study, micro-CT scan analysis revealed that the anterior edge of the swim bladder is physically partitioned by the os suspensorium and that the posterior portion is controlled by the lumen defined by the pleural ribs in adult zebrafish. Also, during the development of the swim bladder, we showed that the os suspensorium on the 4th centrum suppresses the excess inflation of the swim bladder primordium and pushes it back to its proper position. It is known that fish of the carp family including zebrafish have an open swim bladder, in which the airway and swim bladder are connected throughout life (Facey et al., 2022). Thereby, the size of the swim bladder can be regulated by swimming in and out of the swim bladder into and out of the airway. We assume that the mechanism found in our study does not strictly dictate the size and position of the swim bladder, but rather physically restricts the space available for swim bladder inflation to prevent over-expansion. In zebrafish, the swim bladder is known to expand in strong correlation with body growth after the swim bladder is established (Lindsey et al., 2010). In mammals, growth hormones secreted from the pituitary gland are known to play an important role in regulating the growth of essentially all tissues, including the bones (Giustina et al., 2008). The present study suggests the importance of the surrounding skeletons in controlling the size and position of internal organs such as the swim bladder in zebrafish, and that it may play a critical role in the allometric growth of the organs in a tight association of the body growth.

The os suspensorium, which exhibits diverse morphologies in extinct and extant Otophysan fishes (Bird and Hernandez, 2007; Britz et al., 2021; Diogo, 2009), has not been well studied and its role has been unknown. Our results suggest that the zebrafish os suspensorium may play at least two roles. During the inflation of the anterior lobe of the swim bladder, the counterclockwise rotation of the os suspensorium plays a role in pushing back the anterior end of the swim bladder to the proper position. Although the mechanism that drives the movement of the os suspensorium remains unknown, our results suggest that the inflation of the swim bladder in zebrafish is not solely regulated in a cell-autonomous manner, but is rather controlled by the surrounding environment, such as adjacent bones and organs. Second, some studies have mentioned the possibility that the os suspensorium functions as a fifth Weberian ossicle (Bird and Mabee, 2003; Vandewalle et al., 1990), and our results may support this idea. As mentioned above, the os suspensorium has been shown to physically push back the anterior end of the swim bladder during its formation. In addition, the broad contact between the swim bladder and the os suspensorium was observed (Figure 1).

These results suggest that the os suspensorium and the swim bladder were not simply in contact, but the os suspensorium physically pushed back the overinflated swim bladder, suggesting that tension was likely generated between the swim bladder and os suspensorium. The observation that the os suspensorium and swim bladder are in direct contact without a gap in adult zebrafish suggests that this tension is maintained in adult zebrafish. Given that tension is present, even small vibrations from the swim bladder could be more efficiently transmitted to the inner ear. Although the posterior end of the tripus is generally considered to be the primary ossicle for receiving swim bladder vibrations, our results suggest that the os suspensorium, which is in tension with the swim bladder, may be able to more efficiently transmit swim bladder vibrations to the tripus via bone conduction. The four Weberian ossicles show high similarities in morphological structures among Otophysan fishes, whereas the os suspensorium is morphologically diverse among Otophysan fishes, including some Otophysan fishes in which the os suspensorium is underdeveloped (Bird and Hernandez, 2007; Diogo, 2009). Based on these observations, we propose that zebrafish, which have developed the os suspensorium during the Otophysan lineage, may have improved it by physically partitioning the anterior end of the swim bladder and incorporating it into the Weberian apparatus, and have acquired a more efficient mechanism for detecting swim bladder vibrations.

## Supporting information

Satoh et al-Supplementary Data

## Acknowledgments

We thank the National BioResource Project of Japan (zebrafish) for providing the RW strain and the member in our laboratory for the critical discussion. This work was supported by KAKENHI Grants-in-Aid for Scientific Research 18K06177, 23K05790 to AK, NIG-JOINT 38A2019, 7A2020, 66A2021, 18A2022, 31A2023 to AK, and Narishige Zoological Science Award 2021 to AK.

## Author Contributions

K.S., M.A., and A.K. conceived and designed the experiments, isolated *hoxc6a* mutants, K.S., U.A., and K.Y. obtained *hoxc6a* homozygous mutants for micro-CT scan, M.I., R.K., and H.N. obtained *hoxaa* cluster homozygous mutants for micro-CT scan, A.M. performed X-ray micro-CT scan analysis and constructed the 3D-movies, and K.S., U.A., M.I., H.N., S.O., R.F., and A.K. performed the embryonic phenotype analysis. H.F. performed the statistical analysis, and A.K. wrote the manuscript. All authors discussed the results and commented on the manuscript.

## References

Adachi, U., Koita, R., Seto, A., Maeno, A., Ishizu, A., Oikawa, S., Tani, T., Ishizaka, M., Yamada, K., Satoh, K., et al. (2024). Teleost Hox code defines regional identities competent for the formation of dorsal and anal fins. Proc Natl Acad Sci U S A.

Akama, K., Ebata, K., Maeno, A., Taminato, T., Otosaka, S., Gengyo-Ando, K., Nakai, J., Yamasu, K., and Kawamura, A. (2020). Role of somite patterning in the formation of Weberian apparatus and pleural rib in zebrafish. J Anat 236, 622–629.

Alexander, R.M. (1993). Buoyancy (CRC Press).

Bird, N.C., and Hernandez, L.P. (2007). Morphological variation in the Weberian apparatus of Cypriniformes. J Morphol 268, 739–757.

Bird, N.C., and Mabee, P.M. (2003). Developmental morphology of the axial skeleton of the zebrafish, Danio rerio (Ostariophysi: Cyprinidae). Dev Dyn 228, 337–357.

Bird, N.C., Richardson, S.S., and Abels, J.R. (2020). Histological development and integration of the Zebrafish Weberian apparatus. Dev Dyn 249, 998–1017.

Britz, R., Conway, K.W., and Ruber, L. (2021). The emerging vertebrate model species for neurophysiological studies is Danionella cerebrum, new species (Teleostei: Cyprinidae). Sci Rep 11, 18942.

Burton, D., and Burton, M. (2018). Essential Fish Biology.

Cook, V., Groneberg, A.H., Hoffmann, M., Kadobianskyi, M., Veith, J., Schulze, L., Henninger, J., Britz, R., and Judkewitz, B. (2024). Ultrafast sound production mechanism in one of the smallest vertebrates. Proc Natl Acad Sci U S A 121, e2314017121.

Diogo, R. (2009). Origin, Evolution and Homologies of the Weberian Apparatus: A New Insight. Int J Morphol 27, 333–354.

Dong, J., Feldmann, G., Huang, J., Wu, S., Zhang, N., Comerford, S.A., Gayyed, M.F., Anders, R.A., Maitra, A., and Pan, D. (2007). Elucidation of a universal size-control mechanism in Drosophila and mammals. Cell 130, 1120–1133.

Facey, D.E., Bowen, B.W., Collette, B.B., and Helfman, G.S. (2022). The Diversity of Fishes: Biology, Evolution and Ecology, The third edition.

Funk, E.C., Birol, E.B., and McCune, A.R. (2021). Does the bowfin gas bladder represent an intermediate stage during the lung-to-gas bladder evolutionary transition? J Morphol 282, 600–611.

Funk, E.C., Breen, C., Sanketi, B.D., Kurpios, N., and McCune, A. (2020). Changes in Nkx2.1, Sox2, Bmp4, and Bmp16 expression underlying the lung-to-gas bladder evolutionary transition in ray-finned fishes. Evol Dev 22, 384–402.

Giustina, A., Mazziotti, G., and Canalis, E. (2008). Growth hormone, insulin-like growth factors, and the skeleton. Endocr Rev 29, 535–559.

Gokhale, R.H., and Shingleton, A.W. (2015). Size control: the developmental physiology of body and organ size regulation. Wiley Interdiscip Rev Dev Biol 4, 335–356.

Grande, T., and Young, B. (2004). The ontogeny and homology of the Weberian apparatus in the zebrafish Danio rerio (Ostariophysi: Cypriniformes). Zoological Journal of the Linnean Society 140, 241–254.

Korzh, S., Winata, C.L., Zheng, W., Yang, S., Yin, A., Ingham, P., Korzh, V., and Gong, Z. (2011). The interaction of epithelial Ihha and mesenchymal Fgf10 in zebrafish esophageal and swimbladder development. Dev Biol 359, 262–276.

Lindsey, B.W., Smith, F.M., and Croll, R.P. (2010). From inflation to flotation: contribution of the swimbladder to whole-body density and swimming depth during development of the zebrafish (Danio rerio). Zebrafish 7, 85–96.

Maeno, A., Koita, R., Nakazawa, H., Fujii, R., Yamada, K., Oikawa, S., Tani, T., Ishizaka, M., Satoh, K., Ishizu, A., et al. (2024). Hox code responsible for the pattering of the anterior vertebrae in zebrafish. Development in press.

Meeker, N.D., Hutchinson, S.A., Ho, L., and Trede, N.S. (2007). Method for isolation of PCR-ready genomic DNA from zebrafish tissues. BioTechniques 43, 610, 612, 614.

Nelson, J.S., Grande, T.C., and Eilson, M.V.H. (2016). Fishes of the world, The fifth edition.

Parichy, D.M., Elizondo, M.R., Mills, M.G., Gordon, T.N., and Engeszer, R.E. (2009). Normal table of postembryonic zebrafish development: staging by externally visible anatomy of the living fish. Dev Dyn 238, 2975–3015.

Penzo-Mendez, A.I., and Stanger, B.Z. (2015). Organ-Size Regulation in Mammals. Cold Spring Harb Perspect Biol 7, a019240.

Popper, A.N., and Fay, R.R. (1993). Sound detection and processing by fish: critical review and major research questions. Brain Behav Evol 41, 14–38.

Robertson, G.N., McGee, C.A., Dumbarton, T.C., Croll, R.P., and Smith, F.M. (2007). Development of the swimbladder and its innervation in the zebrafish, Danio rerio. J Morphol 268, 967–985.

Rosello-Diez, A., and Joyner, A.L. (2015). Regulation of Long Bone Growth in Vertebrates; It Is Time to Catch Up. Endocr Rev 36, 646–680.

Rosen, D.E., and Greenwood, P.H. (1970). Origin of the Weberian apparatus and the relationships of the ostariophysan and gonorynchiform fishes.. American Museum Novitates 2428, 1–25.

Seymour, R.S., Christian, K., Bennett, M.B., Baldwin, J., Wells, R.M., and Baudinette, R.V. (2004). Partitioning of respiration between the gills and air-breathing organ in response to aquatic hypoxia and exercise in the pacific tarpon, Megalops cyprinoides. Physiol Biochem Zool 77, 760–767.

Texada, M.J., Koyama, T., and Rewitz, K. (2020). Regulation of Body Size and Growth Control. Genetics 216, 269–313.

Tumaneng, K., Russell, R.C., and Guan, K.L. (2012). Organ size control by Hippo and TOR pathways. Curr Biol 22, R368–379.

Vandewalle, P., Radermaker, F., Surlemont, C., and Chardon, M. (1990). Apparition of the Weberian characters in Barbus barbus Linnaeus 1758 Pisces Cyprinidae. Zoologischer Anzeiger 225, 362–376.

Wallace, K.N., and Pack, M. (2003). Unique and conserved aspects of gut development in zebrafish. Dev Biol 255, 12–29.

Yamada, K., Maeno, A., Araki, S., Kikuchi, M., Suzuki, M., Ishizaka, M., Satoh, K., Akama, K., Kawabe, Y., Suzuki, K., et al. (2021). An atlas of seven zebrafish hox cluster mutants provides insights into sub/neofunctionalization of vertebrate Hox clusters. Development 148.

Zheng, W., Wang, Z., Collins, J.E., Andrews, R.M., Stemple, D., and Gong, Z. (2011). Comparative transcriptome analyses indicate molecular homology of zebrafish swimbladder and mammalian lung. PLoS One 6, e24019.

