## Supplementary material for "Physical constraints on the positions and dimensions of the zebrafish swim bladder by surrounding bones": Satoh et al-Supplementary Data

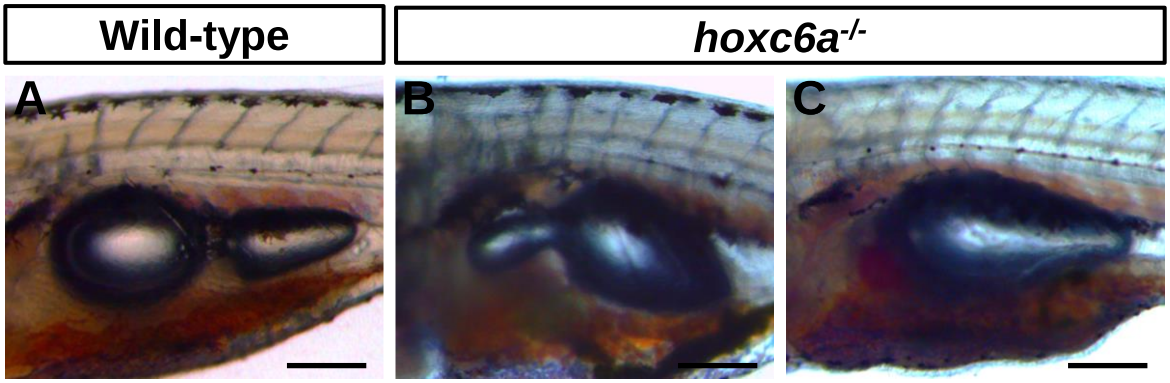


**D**

| | *n* | Normal | Small size of the anterior lobe | Absence of the anterior lobe |
|---|---|---|---|---|
| Wild-type | 48 | 46 (95.8%) | 2 (4.2%) | 0 (0%) |
| *hoxc6a*$^{-/-}$ | 60 | 0 (0%) | 32 (53.3%) | 28 (46.7%) |

**FIGURE S1**

Zebrafish *hoxc6a* mutants exhibit defects in the formation of the anterior lobe of the swim bladder. (A-C) Representative images of the swim bladder in wild-type and *hoxc6a* homozygous larvae . Lateral view. Scale bar: 200 μm. (D) Comparisons of the morphology of the swim bladder between wild-type and *hoxc6a* mutants. As *hoxc6a* homozygous fish demonstrate reproductive capability, larvae obtained from the intercrosses between *hoxc6* homozygous fish were used in this analysis. We have confirmed that phenotype of maternal-zygotic *hoxc6a* mutants is indistinguishable from zygotic *hoxc6a* homozygous fish.
